# ThirdPeak: A flexible tool designed for the robust analysis of two- and three-dimensional (single-molecule) tracking data

**DOI:** 10.1101/2024.01.23.576823

**Authors:** Thomas Müller, Elisabeth Meiser, Markus Engstler

## Abstract

ThirdPeak is an open-source tool tailored for the comprehensive analysis of two– and three-dimensional track data across various scales. Its versatile import options facilitate seamless integration into established workflows, and the user interface allows for swift visualization and analysis of the data. When applied to live-cell diffusion data, this software unveils the advantages of combining both 2D and 3D analysis, providing valuable insights into the understanding of biological processes.

## Main

A wide array of molecules and systems has been probed by single molecule localization microscopy (SMLM). Continuous enhancements in fluorescent dyes ^1^, camera sensors ^2^ as well as localization and tracking algorithms ^3,4^ contribute to the ongoing evolution of this field.

While many processes are readily observable in two dimensions, exploring the third dimension in single molecule microscopy presents a more intricate challenge, often associated with substantial requirements in optical hardware. An effective technique for determining the axial position of individual particles in SMLM involves modifying the point spread function (PSF), e.g. by adding an astigmatic lens into the emission path, to encode the emitter’s position relative to the focal plane ^5^.

Following the acquisition of image data, automating the localization procedure of the emitters can be accomplished through freely available software such as SMAP^6^ or Picasso^7^. Subsequently, the localized points can be connected into tracks, ultimately unveiling the behavior of the target molecules. Several software tools, including TrackMate^8^, uTrack^9^, Swift^10^, or Tardis^4^, serve this specific purpose.

Although existing tools can extract specific information from these tracks, the implementation of interactive data exploration is often lacking. This aspect, however, becomes crucial in the context of biological data, where omnipresent noise can obscure important characteristics. While interactive tools for two-dimensional data do exist, they are constrained by their special data structure. Although a standard for SMLM data^12^ has been proposed, its general adoption is still pending. Therefore, a notable gap currently exists in open and user-friendly applications for exploring and analyzing three-dimensional data (Sup. Fig. 1).

Here, we introduce ThirdPeak, a software designed to visualize, and analyze three-dimensional track data with a focus, but certainly not limited to, SMLM. The graphical user interface ensures accessibility for a diverse range of scientists, while the versatile data import feature facilitates integration of the software into established workflows.

To ensure seamless integration with both existing and future workflows, the software operates with established formats and accommodates custom MATLAB and comma-separated value (CSV) files via an import dialogue. This flexibility enables the analysis of data spanning a variety of spatial and temporal scales (Fig. 1 A, Sup. Fig. 2).

**Figure 1:**
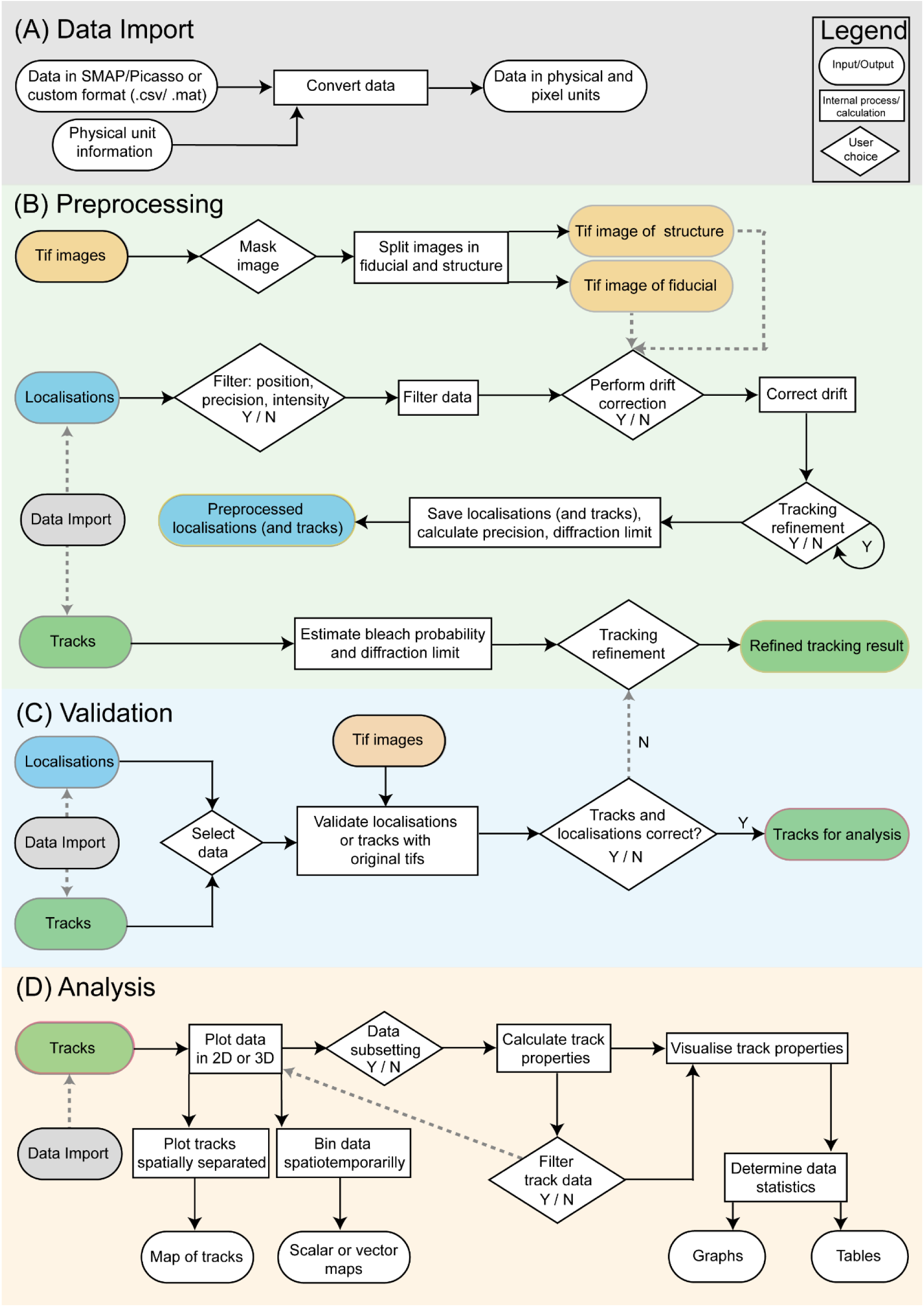
Data flow for the data import, preprocessing, validation and analysis in ThirdPeak. Illustrating the input and output steps (ellipses), the choices for the user (diamond) and the internal processing of the software (rectangle). Y/N: Yes/No.

The preprocessing stage (Figure 1 B, Sup. Fig. 2) enables batch-processing of single-molecule localizations. First, data can be filtered based on position, localization precision, and intensity. To mitigate the prevalent issue of drift in small-scale data, drift corrections can be applied. With the installation of Swift^10^ on the system, tracking of localizations becomes seamlessly achievable. Alternatively, parameters such as the distribution of localization precision and the diffraction limit are saved and serve as input parameters for a preferred tracking algorithm.

Using track data facilitates the computation of estimated displacement and bleaching probability of the particles. These values subsequently contribute to refining the tracking process iteratively.

During the validation phase (Figure 1 C), users can interactively navigate through both localizations and tracks. This data can be overlaid onto the original microscopy images, facilitating the identification of any necessary adjustments in either the localization or tracking steps. Subsequently, users can proceed to the visualization window for a comprehensive data analysis (Figure 1 D).

The central window offers the flexibility to manipulate the three-dimensional data, enabling exploration from various viewpoints. On the right-hand side of the visualization window, users have the option to manually subset the data and save the selections for later analysis (Sup. Fig. 2).

The analysis aims to derive standard parameters, including jump distances and diffusion coefficients. The distribution of the mean jump distance is particularly valuable for identifying distinct diffusing populations.

A data filtering feature is provided for all calculated properties, enabling users to further narrow down the data based on their interests. This feature helps in uncovering concealed dynamics within selected groups. These data filter settings can be saved and loaded, thereby enhancing the reproducibility of the data analysis (Sup. Fig. 2).

The entirety of the data can be exported in CSV or Excel formats. Moreover, the generated figures can be exported as a scalable vector graphic, suitable for presentations and publications. This streamlined process provides a swift and accessible method for exploring and comprehensively analyzing three-dimensional track data, eliminating the necessity for programming expertise.

To demonstrate the software’s capabilities, artificial diffusion data of three mixed populations was generated using SMIS^13^. In addition, experimental data was acquired from living, immobilized *Trypanosoma brucei,* using astigmatism to encode the axial position (Sup. Fig. 3). Single molecules were localized using SMAP, followed by preprocessing of the localization data in ThirdPeak. The filtered data was tracked using swift and subsequently visualized and analyzed using ThirdPeak again (Fig. 2, Sup. Fig. 4).

**Figure 2:**
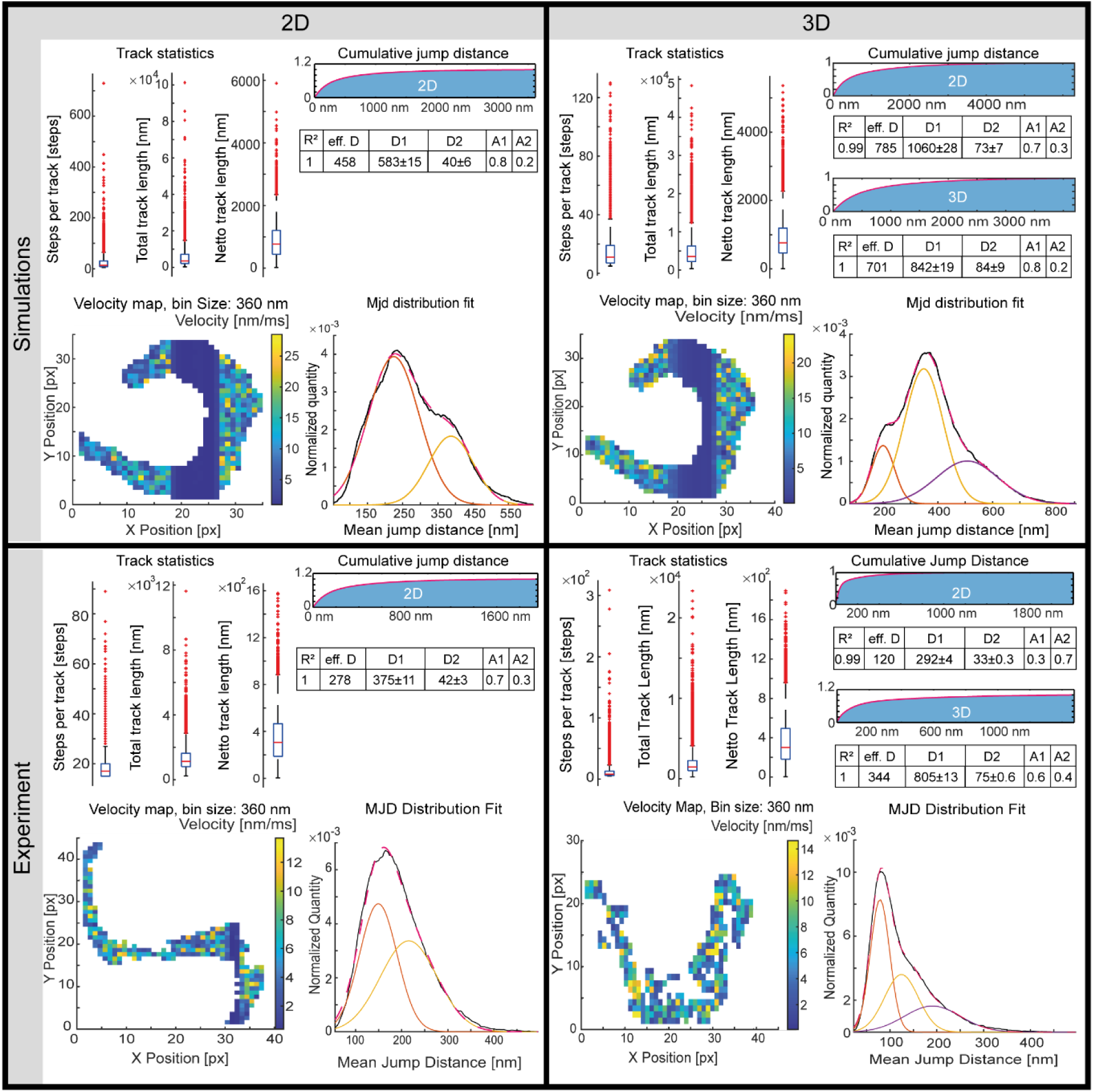
Simulated and experimental diffusion data of the surface coat of *Trypanosoma brucei*. Overall track statistics, a fit to the cumulative jump distance distribution to determine the diffusion coefficients, a velocity map and the mean jump distance distribution are shown.

A comparison of track statistics between the simulated and experimental datasets reveals only minor differences in the number of steps tracked and the total track length. Notably, when examining the net track length (distance between the first and last localization), the experimental data exhibits lower values compared to the simulation. This observation implies the potential presence of either confinement or fast bleaching processes that are not present in the “ideal” simulation (Sup. Table 1). The diffusion coefficients through fitting to the cumulative squared jump distance distribution and are in good agreement with previously determined 2D diffusion data^14^(Sup. Fig. 4, Sup. Table 1). The binning in velocity maps serves to reduce noise by spatial averaging and reveal an unusually slow region in the center of the simulated data. In the experimental data, an increased velocity is frequently observed along the groove of the attached flagella. The mean jump distance analysis can reveal either two or three diffusing populations, depending on the dimensionality of the data.

To the best of our knowledge, ThirdPeak is the first software to seamlessly integrate interchangeable 2D and 3D track analysis. We anticipate that ThirdPeak will prove beneficial for users of different backgrounds. Our commitment includes ongoing efforts to improve the software by incorporating additional features in future updates.

## Acknowledgments

We would like to thank Pierre Parutto (Cambridge, UK) and Daniele Bourgeois (Grenoble) for sharing their work and helpful discussion. We further would like to thank Philip Kollmannsberger (Düsseldorf) and Sabine Fischer (Würzburg) for discussion and advice.

## Supplementary text and figures

**Supplementary figure 1:**
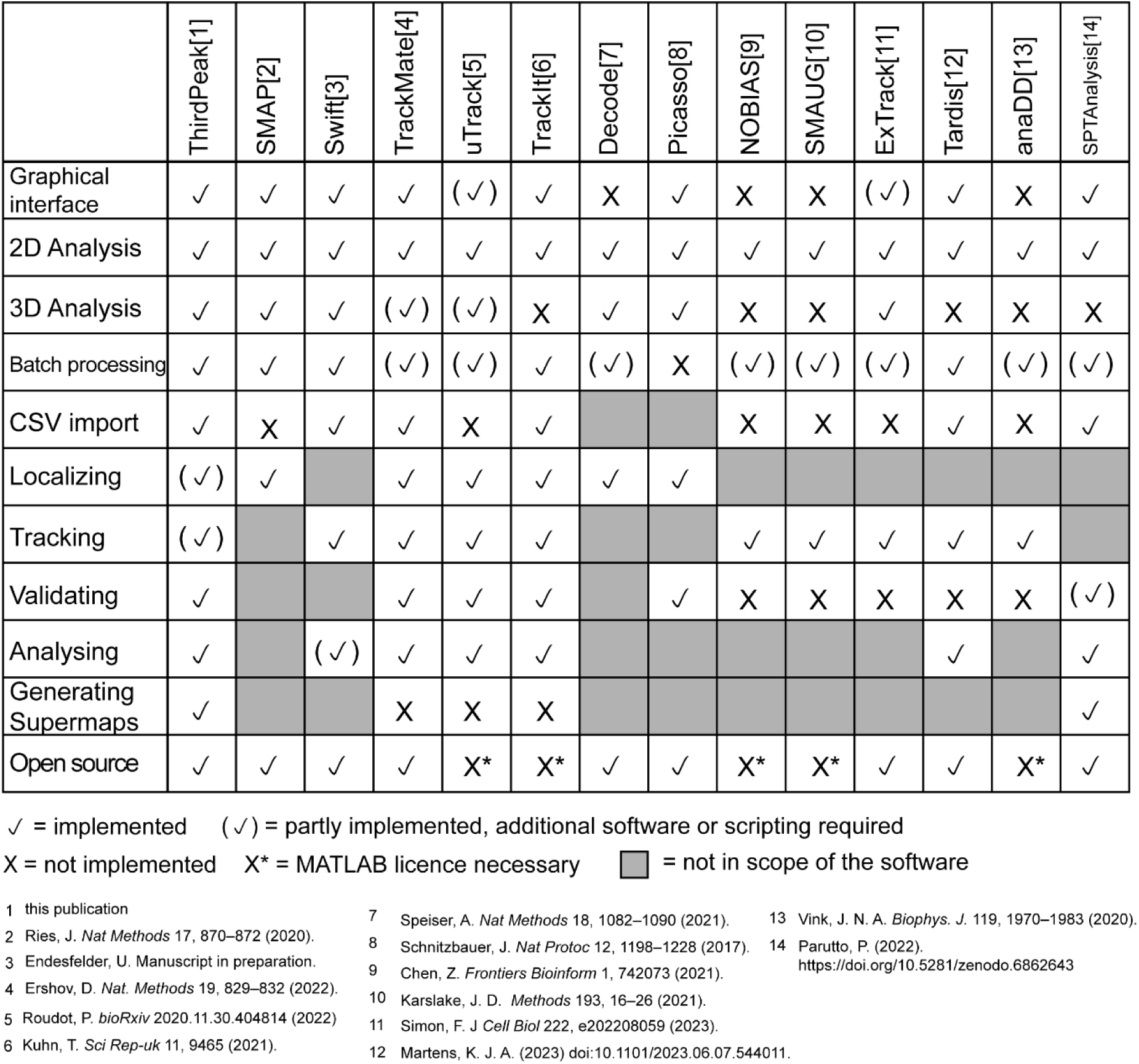
**Overview of software for single molecule localization and tracking**.

**Supplementary figure 2:**
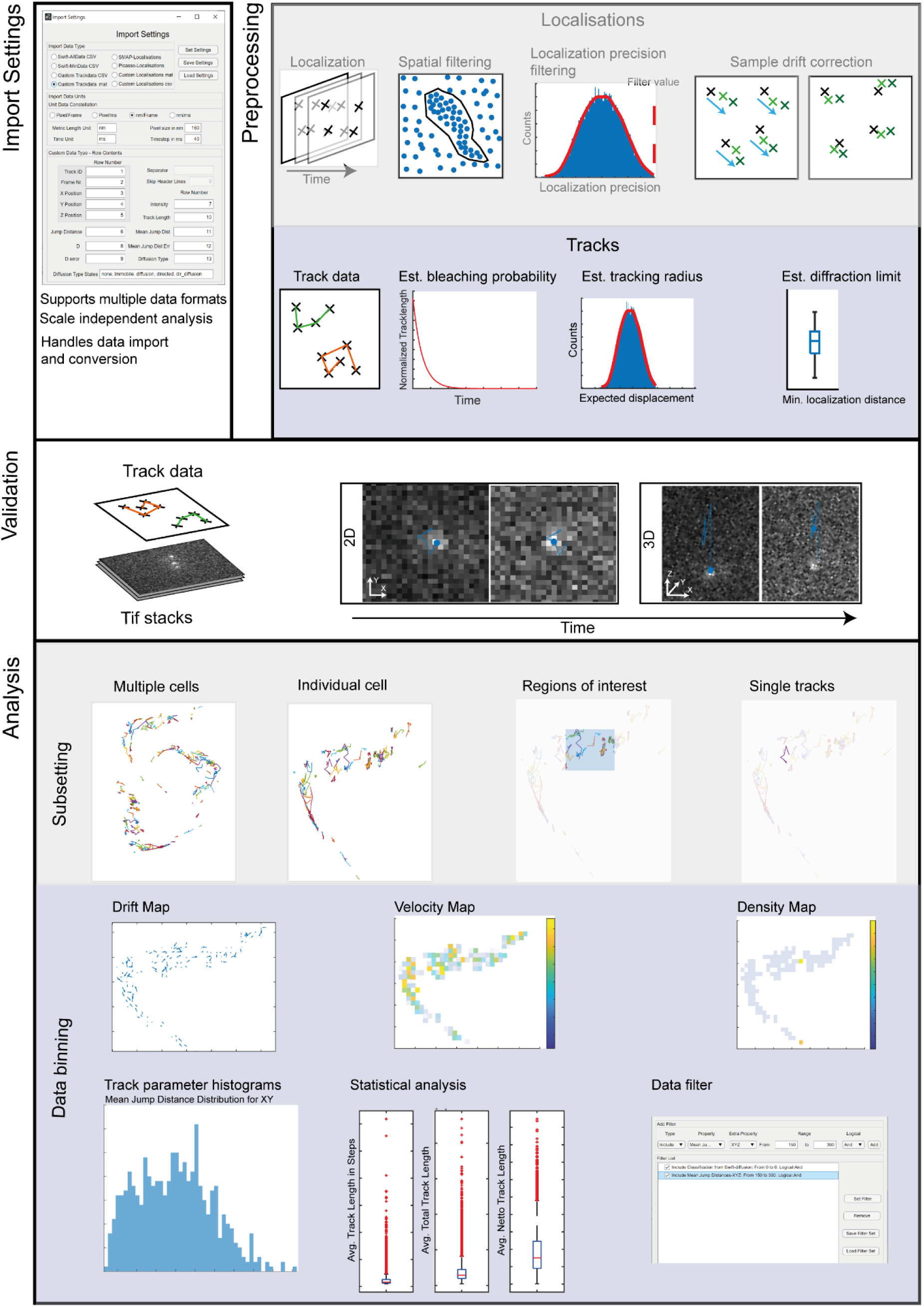
Feature overview of ThidPeak on an exemplaray 3D data set of surface diffusion on *Trypanosoma brucei*. The import dialogue needs to be set in the beginning of the chosen workflow (preprocessing, validation or analaysis) to determine the expected data format. During preprocessing, localisation data can be filtered on their spatial position, their precision values or their intesity. Histograms of the remaining localisations are generated and saved in the current working directory during the processing. An automatic drift correction can be applied if desired to the data, either using a mean shift approach or fiducials. The processed localisations can then be tracked if swift is installed on the computer directly from the GUI of ThirdPeak. Futher, generated track data can be loaded into the preprocessing workflow to determine bleach probability, diffraction limit and the expected displacement value to refine the tracking process. If enough data is present, one can also calculate the fourier ring correlation or fourier shell correlation to determine the resolution of the resolved structure. If only a few houndred data points are present, the diffraction limit can be determined by the minimal distance of the localisation per frame. During the validaiton step, localisaiton or track data can be used with the original tif images to check and, if necessary, refine the localisation and tracking results. The analysis workfow allows the visualisation of the track data, subsetting it into single tracks or region of interest. Alternatively, the data can be binned to visualize locally dominating dynamics, either by their overall drift or their velocities. Data can then be visualized in histograms or used for statistical analysis. The track data can then be further filtered using the calculated track properties and evaluated again.

**Supplementary figure 3:**
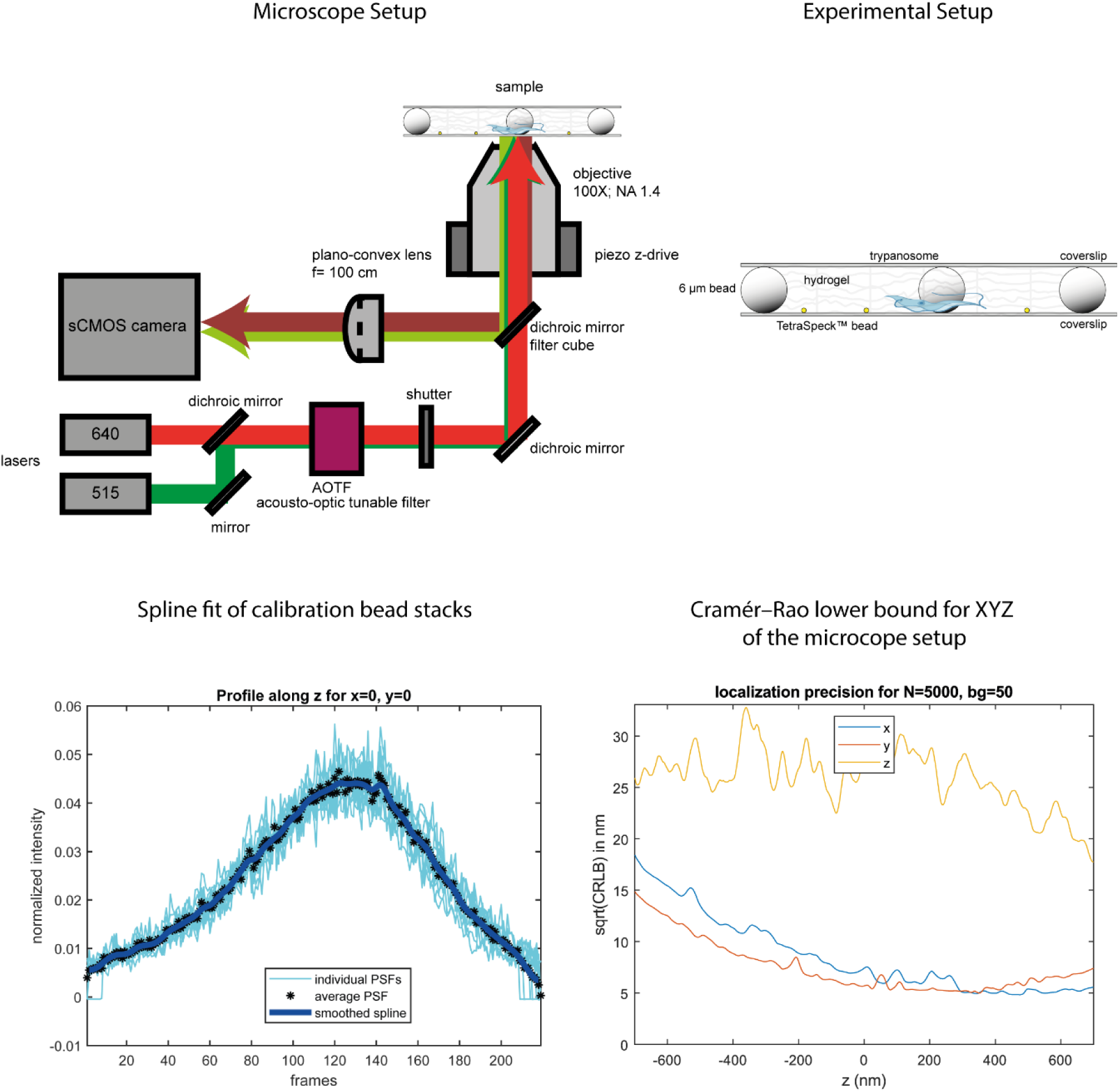
Microscope and experimental setup used for the experimental data. Calibration and Cramer-Rao lower bound of the astimgatic setup. For single molecule imaging, a standart widefield microscope is used. We use two laser lines at 515 nm and 640 nm respectively. The protein of interest is labelled with JF646 and illuminated by the 640 nm laser. The cytosol is labelled with orgeon green for finding and focusing on the cells. The laser line can be quickly switched using an AOTF. The emitted light from the fluorophores is then directed through the astigmatic lens to encode the z position of the emitters by the shape of their point spread function. To image living, immobilized cells we use a hydrogel based on polyethyleneglycole and hyaluronic acid. The cells are resuspended in the gel and placed on a clean coverslip. Furthermore, spacer beads are included to prevent squeezing of the cells in between the two coverslips. For the determination of the z position of the single emitters, calibration images of fluorescent TetraSpek beads in the beforemention hydrogel are acquired. With these image stacks, a calibration curve can be determined by SMAP. From this calibartion curve, the calculated Cramer-Rao lower bound can be determined, which describes the therotically best possible precision for the given dimension. Due to the astigamtism and the associated z-dependend shape change of the point spread function, the X and Y precision is diminished at the lower and upper limit, with the best precision at the center. For z, the resolution is always worse then for X and Y and is usually around 35-40 nm in the calibrated range.

**Supplementary figure 4:**
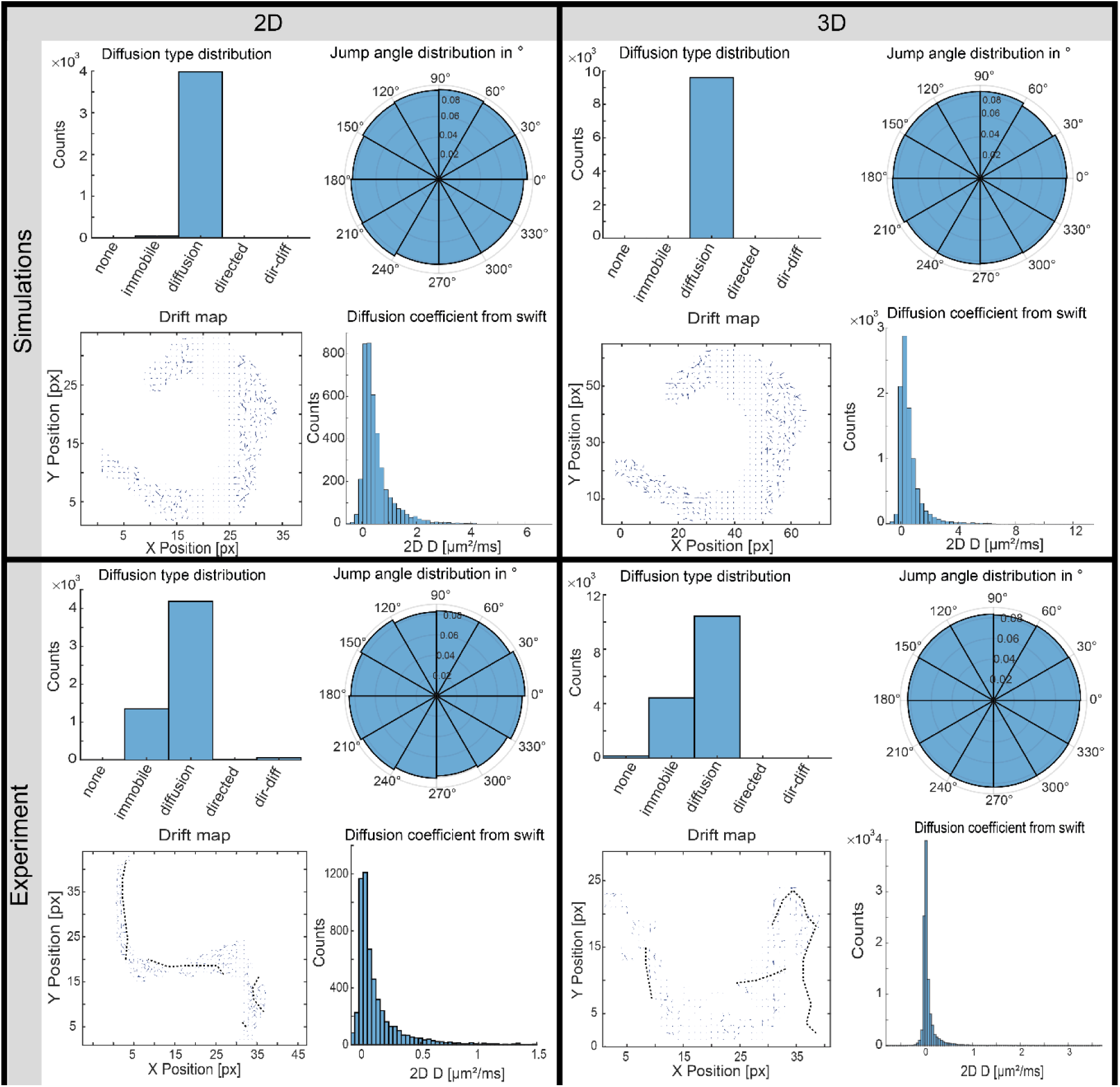
Additional track properties of simulated and experimental diffusion data. The simulated data shows that all tracked particles were diffusive, while for the experimental data, a smaller fraction of 20-30% was considered immobile by the swift algorithm. A particle is considered immobile if it is not moving more than the value determined by the localisation precision. The jump angle distribution shows an even distirbution for all conditions. A primary jump angle around 0° could point to directed transport, while a jump angle of 180° could indicate a large amount of immobile particles. The drift map illustrates the overall direction of the tracks on their given position in the grid. For the simulations, neither sample drift nor directionality onto a given position is visible. In the center part of the simulated area, the drift vectors are small, coinciding with the small velocity values, which is probably a result of the model used. For the experimental data, drift vectors are often paralell to the flagella, as particles can not usually cross the flagellar membrane. A circle of drift vectors pointing all to the center of the circle can indicate the flagellar pocket of the cell, which is not present in the simulated data. The diffusion coefficient determined by swift does not show a large difference between the data sets. It has to be noted that this diffusion coefficient is only determined in 2D and not for all three dimensions.

**Supplementary Table 1:**
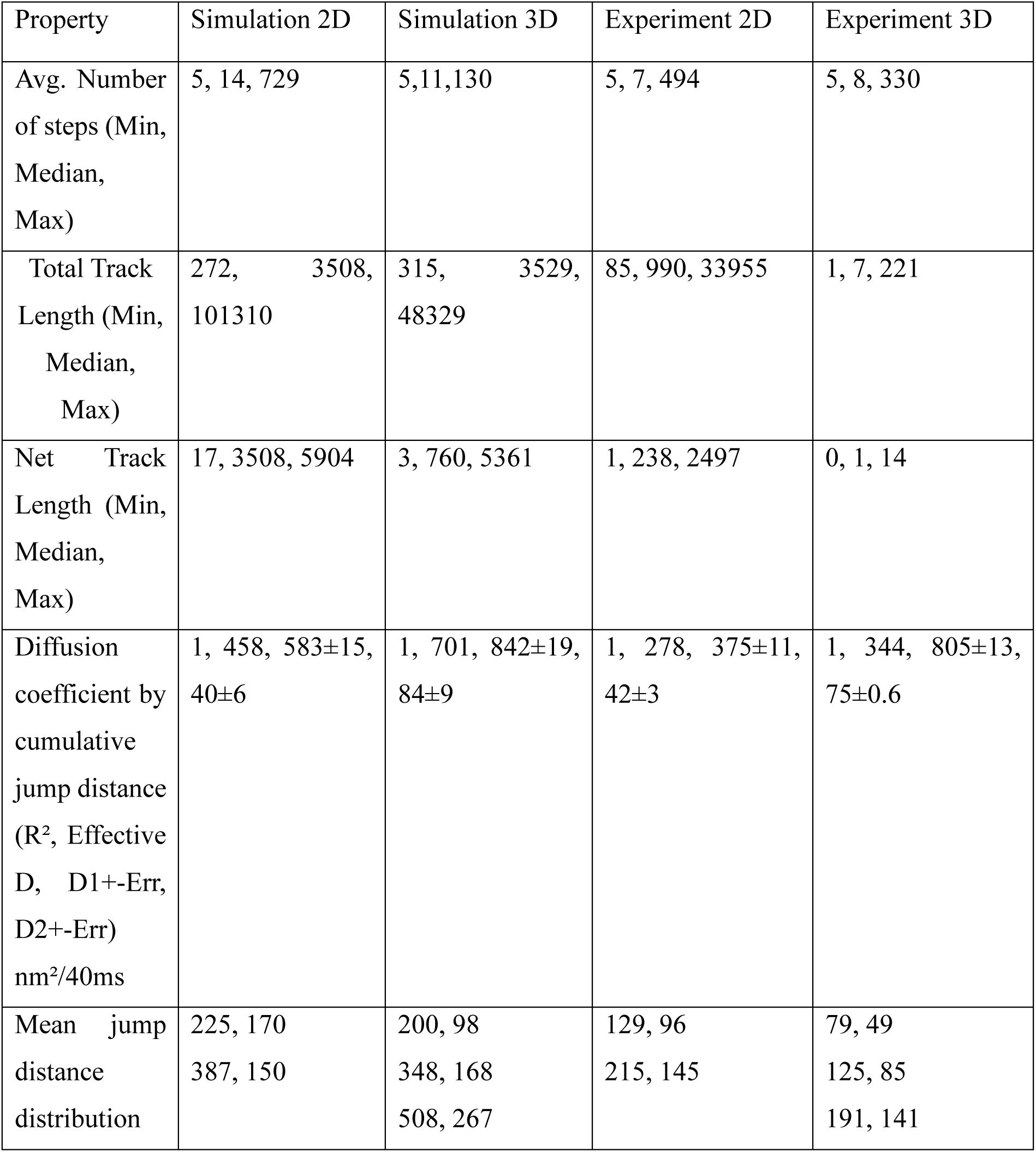
Results of the track analysis.

**Supplementary figure 5:**
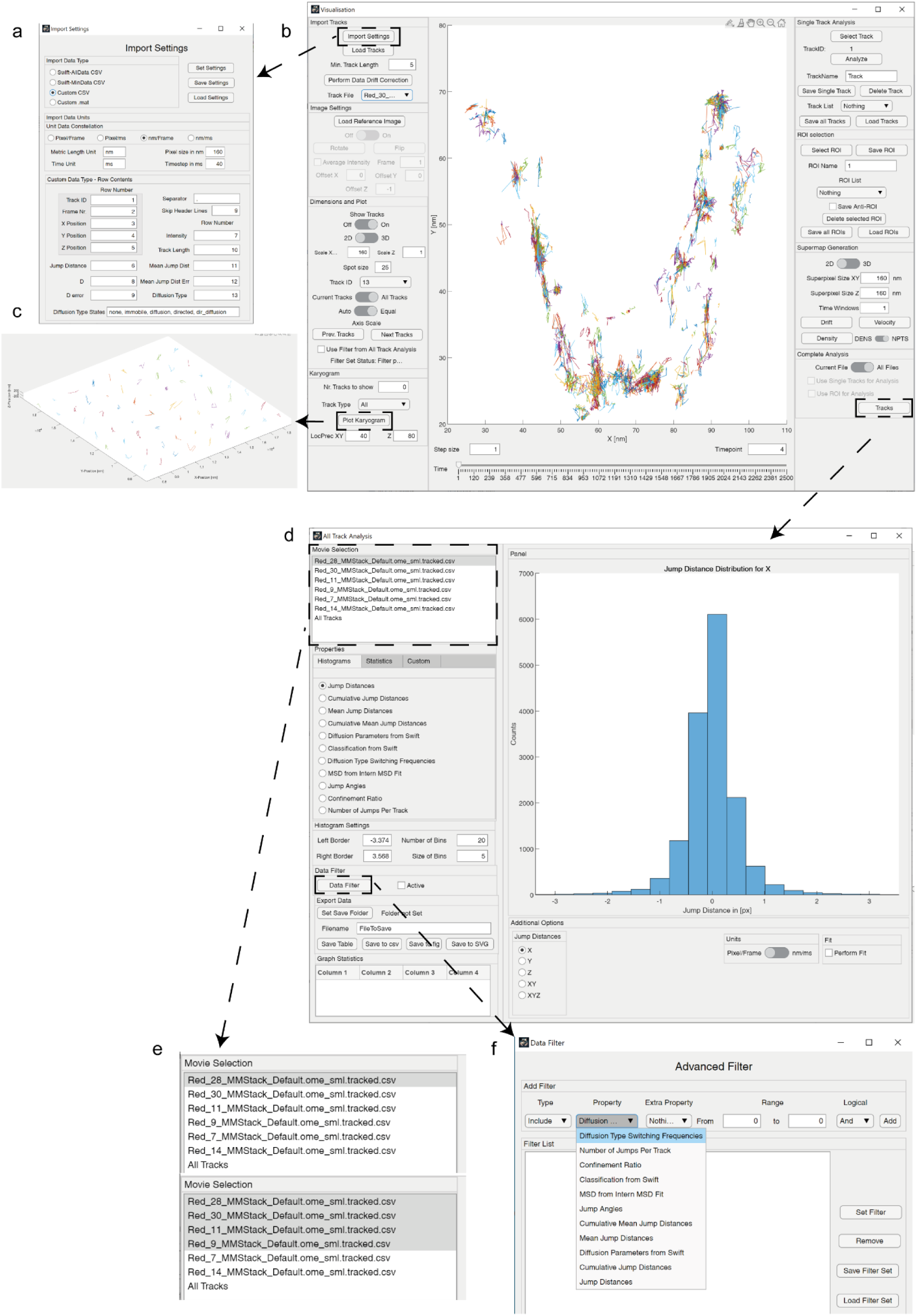
Screenshots of ThirdPeak. A shows the Import dialogue for the data and scaling of the data. B shows the general overview of the visualisation workflow. C an overview of the tracks, lined up in a grid. D depicts the analysis window right after opening it. E allows the selection of the data, either a single file, multiple files or all files can be selected. F shows the options of the advanced filter dialogue in more detail.

## Methods

The software was developed using MATLAB 2021 and 2022 using Windows 10.

## Simulations

The simulation of the single molecule data was done using SMIS2.1, running in MATLAB 2022 in Windows 10. Imaging parameters were chosen to represent the local single molecule microscope.

## Cell culture and cell line generation

Trypanosoma brucei strain Lister 427 were cultivated at 37°C, 5% CO2 in HMI-9 (*Hirumi’s Modified* Iscove’s medium-*9)* containing 10% heat-inactivated fetal bovine serum.

## Sample preparation and imaging

Thickness-corrected coverslips were placed in coverslip-racks made from Teflon and cleaned two times using 2% Hellmanex-II solution by incubating them for ten minutes in a ultrasonic water bath (Elmasonic P, Elmasonic) using 100% power, a frequency of 37 kHz at room temperature. Coverslips were then washed using ddH2O and stored in ddH2O until use. On the day of the experiment, two coverslips per sample were dried using an air stream and placed in a clean petri dish. For one experimental day, 1×10^7 cells were harvested by centrifugation at 1500 xg for ten minutes at RT. The supernatant was removed, and the cells resuspended in one ml of vPBS (PBS supplemented with 10 mM glucose and 46 mM sucrose, pH 7.6). Cells were washed three times using one ml of vPBS and subsequent centrifugation at 800 xg for two minutes. After the last washing step, the cells were resuspended in 100 µl vPBS containing 1 nM JaneliaFluor646-NHS and incubated for five minutes on ice. The dye would then label the homogenous surface coat consisting of variant surface glycoproteins (VSG). After the incubation period, the cells were washed three times with 1 ml of vPBS, again centrifuged at 800 xg for two minutes at 4°C. At the end, the cells were resuspended in 20µl vPBS and stored on ice. As the cells are flagellated unicellular parasites, they had to be immobilized for imaging. For this, a mixture of 8-arm PEG-VS (2.5 µl of 50 mM solution) is used together with a hyaluronic-acid that was functionalized with an SH group (8-10 kDa, DS 40%, 3 µl of 25% solution). Additionally, beads of six µm diameter were used as a spacer, 2 µl vPBS as a buffer and 2 µl of the cell solution. The mixture was well mixed and added onto a coverslip. A second coverslip was placed on top to seal the sample and a weight of 90 g was applied to the sample to make sure that it is even. The sample was then centrifuged for 1 min at 1500 xg to make sure that most of the cells were plain on the glass at the bottom coverslip.

Samples were imaged at 37°C using a heating chamber on a Leica microscope with a oil immersion objective 100×1.4 numerical aperture (Olympus). The sample was excited using a laser at 640 nm with a power of 3 kW/cm². Pulsed illumination was controlled using an acusto-optical tunable filter AOTF to reach illumination times of 9 ms or 36 ms at frame rates of 100 Hz or 25 Hz for 2D and 3D respectively. Either 5000 or 2500 frames were acquired. Image acquisition was performed using an Andor iXon Ultra with a gain of 150, running in cropped mode with a ROI of 120×120 pixel. The acquisition process was controlled using MicroManager 2.0.

For the acquisition of calibration samples for the 3D astigmatic approach, TetraSpeck beads were immobilized using the PEG-HASH hydrogel and z-stacks with 10 nm step size were acquired.

Image processing

Calibration stacks of fluorescent beads were processed in SMAP to determine the astigmatism using a spline fit. This generated calibration file could then be used for the z-position determination of the emitters on the actual sample. For this, the batch-mode of SMAP was used. The localizations were then loaded into ThirdPeak, and a manual mask was applied to only include data close to the cell shape. Further, localizations were filtered by their precision (XY max 100 nm, Z max. 200 nm). After that, localizations were corrected for drift using a mean-shift approach and remaining localizations were saved as CSV files to be analyzed by swift. After tracking via swift, the tracking results were loaded into ThirdPeak to refine the tracking by adjusting the expected displacement and bleach values accordingly. After the tracking process converges to a certain expected displacement, the tracking process was optimized. These optimized tracks were then loaded into ThirdPeak for further analysis.

## Functions in the software

### Supermap generation

As biological data is inherently noisy, generating supermaps by binning single-molecule localizations and tracks into larger areas can reveal processes that might just be hidden inside the noise itself. For this, the data range of the tracks is used, and the x, y and z coordinates are placed into respective bins, depending on the size chosen by the user. Additionally, one can choose to also change the temporal binning by increasing the number of time windows, thereby splitting the data also in the temporal space. This feature is available for the drift of the tracks, which could reveal an overall direction of the particles in relation to their positioning, the velocity of particles and the number of localizations in a given bin, either as the absolute number or the relation between the number and the bin size.

### Jump Distance

The jump distance describes the Euclidean distance of a particle between two consecutive timepoints in 1D, 2D or 3D, depending on the choice of the user. For one dimension:

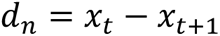

Where d describes the distance between two time points in one of the dimensions, xt the positional argument of one dimension at timepoint t and xt+1 the positional argument of one dimension at timepoint t+1.

For two dimensions:

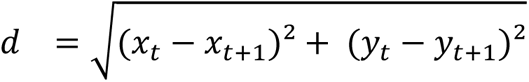

And three dimensions:

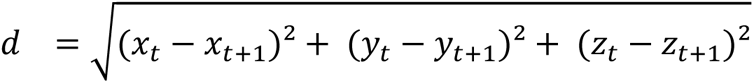

### Mean Jump Distance

The mean jump distance can be calculated per track from the jump distances calculated above by:

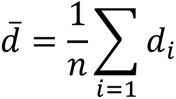

### Net Track Distance

The net track distance describes the distance between the first and last localization of a particle,

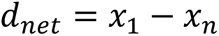

### Total Track Distance

The total track distance describes the complete length of a trajectory.

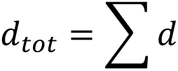

### Confinement ratio

The confinement ratio divides the total track length by the net distance. If this value is close to 1, the track is not confined. If it is much larger than 1, the track is very confined.

### Diffusion coefficient estimation by Jump Distance Distribution

Estimating the diffusion coefficient from the 1D jump distance distribution provides are robust way of calculating this property under the assumption of Brownian motion, especially when having short tracks, as a fit to the mean-squared displacement becomes less reliable. To retrieve the diffusion coefficient, fitting a normal distribution onto the jump distance distribution of one dimension can be performed. The standard deviation from this fit can then be retrieved and by the Einstein-Smoluchowski equation the diffusion coefficient can be calculated by

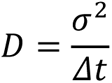

Where D is the diffusion coefficient, σ is the standard deviation and Δ*t* the respective timeframe. This however only determines the overall diffusion coefficient and can not distinguish between different populations.

### Diffusion coefficient estimation by Cumulative Jump Distance Distribution

To distinguish between multiple populations, an analysis using the cumulative jump distance distribution can be performed by fitting the integrated distribution in accordance to Weimann, Klenerman for 2D

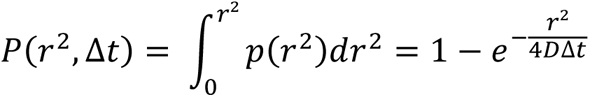

For one species and if multiple species are present, the sum is used:

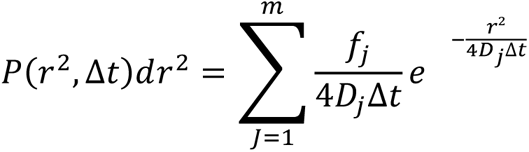

And for 3D

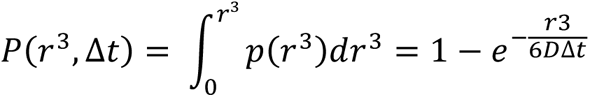

### Mean jump distance population distribution

The mean jump distance population distribution is determined by taking the histogram of the mean jump distances, smoothing it with a variable kernel filter and determining the maxima in of the smoothed data. Those determined maxima are then used in a minimal least squared approach to fit the data with gaussian populations.

### Confinement Ratio vs mean jump distance

The confinement ratio of a particle can be determined by fitting the mean squared displacement with a confined diffusion model:

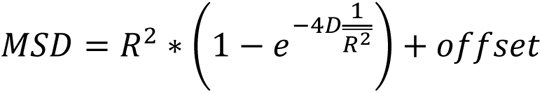

Where R is the radius of confinement and D the local diffusion coefficient.

### Volume calculations

Volume calculations are performed on the total volume the tracks confine. Two options are available, the convex hull, that forms a convex bounding box around the track data, while the alphaShape approach will fit a tight bounding box around the track data.

### Jump Angle

The angle δ between two vectors, or essentially three points of a single track can be calculated as:

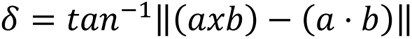

With a and b being the two vectors respectively.

## Data availability

Data related to this article are available on:

## Code availability

The source code as well as compiled versions as a MATLAB app and standalone executable for Windows and MacOS are available on Github: https://github.com/unithmueller/ThirdPeak

## References

1. Grimm, J. B. et al. A General Method to Improve Fluorophores Using Deuterated Auxochromes. JACS Au 1, 690–696 (2021).

2. Fujiwara, T. K. et al. Development of ultrafast camera-based single fluorescent-molecule imaging for cell biology. J. Cell Biology 222, e202110160 (2023).

3. Speiser, A. et al. Deep learning enables fast and dense single-molecule localization with high accuracy. Nat Methods 18, 1082–1090 (2021).

4. Martens, K. J. A., Turkowyd, B., Hohlbein, J. & Endesfelder, U. Temporal analysis of relative distances (TARDIS) is a robust, parameter-free alternative to single-particle tracking. (2023) doi:10.1101/2023.06.07.544011.

5. Diezmann, A. von, Shechtman, Y. & Moerner, W. E. Three-Dimensional Localization of Single Molecules for Super-Resolution Imaging and Single-Particle Tracking. Chem Rev 117, 7244–7275 (2017).

6. Ries, J. SMAP: a modular super-resolution microscopy analysis platform for SMLM data. Nat Methods 17, 870–872 (2020).

7. Schnitzbauer, J., Strauss, M. T., Schlichthaerle, T., Schueder, F. & Jungmann, R. Super-resolution microscopy with DNA-PAINT. Nat Protoc 12, 1198–1228 (2017).

8. Ershov, D. et al. TrackMate 7: integrating state-of-the-art segmentation algorithms into tracking pipelines. Nat. Methods 19, 829–832 (2022).

9. Roudot, P., et al. u-track 3D: measuring and interrogating dense particle dynamics in three dimensions. bioRxiv 2020.11.30.404814 (2022) doi:10.1101/2020.11.30.404814.

10. Endesfelder, M., Schießl, C., Turkowyd, B., Lechner, T. & Endesfelder, U. swift – fast, probabilistic tracking for dense, highly dynamic single-molecule data. manuscript in prep.

11. Kuhn, T., Hettich, J., Davtyan, R. & Gebhardt, J. C. M. Single molecule tracking and analysis framework including theory-predicted parameter settings. Sci Rep-uk 11, 9465 (2021).

12. Sage, D. et al. Super-resolution fight club: assessment of 2D and 3D single-molecule localization microscopy software. Nat Methods 16, 387–395 (2019).

13. Bourgeois, D. Single molecule imaging simulations with advanced fluorophore photophysics. Commun Biology 6, 53 (2023).

14. Schwebs, M. et al. Single-molecule fluorescence microscopy demonstrates fast dynamics of the variant surface glycoprotein coat on living trypanosomes. Biorxiv 2022.08.03.502583 (2022) doi:10.1101/2022.08.03.502583.

